# Exploring the role of the metabolite-sensing receptor GPR109a in diabetic nephropathy

**DOI:** 10.1101/713354

**Authors:** Matthew Snelson, Sih Min Tan, Gavin C. Higgins, Runa Lindblom, Melinda T. Coughlan

## Abstract

Alterations in gut homeostasis may contribute to the progression of diabetic nephropathy. There has been recent attention on the renoprotective effects of metabolite-sensing receptors in chronic renal injury, including the G-protein-coupled-receptor (GPR)109a, which ligates the short chain fatty acid butyrate. However, the role of GPR109a in the development of diabetic nephropathy, a milieu of diminished microbiome-derived metabolites, has not yet been determined. This study aimed to assess the effects of insufficient GPR109a signalling via genetic deletion of GPR109a on the development of renal injury in diabetic nephropathy. *Gpr109a*−/− mice or their wildtype littermates (*Gpr109a*+/+) were rendered diabetic with streptozotocin (STZ). Mice received a control diet or an isocaloric high fiber diet (12.5% resistant starch) for 24 weeks and gastrointestinal permeability and renal injury were determined. Diabetes was associated with increased albuminuria, glomerulosclerosis and inflammation. In comparison, *Gpr109a*−/− mice with diabetes did not show an altered renal phenotype. Resistant starch supplementation did not afford protection from renal injury in diabetic nephropathy. Whilst diabetes was associated with alterations in intestinal morphology, intestinal permeability assessed *in vivo* using the FITC-dextran test was unaltered. GPR109a deletion did not worsen gastrointestinal permeability. Further, 12.5% resistant starch supplementation, at physiological concentrations, had no effect on intestinal permeability or morphology. These studies indicate that GPR109a does not play a critical role in intestinal homeostasis in a model of type 1 diabetes or in the development of diabetic nephropathy.

## Introduction

Diabetic nephropathy is a major microvascular complication of diabetes, occurring in up to 30% of patients with type 1 diabetes [1]. Concomitant with the rise in diabetes and obesity, the prevalence of diabetic nephropathy has been increasing rapidly, with diabetic nephropathy now the leading cause of end stage renal disease worldwide [1]. Despite optimal conventional management with pharmacological inhibition of the renin angiotensin system (RAS), glycemic and blood pressure control, a significant proportion of patients with DKD still progress over time to end stage renal failure. Thus, there is an urgent need for the identification of new therapeutic options to help limit the progression of this disease.

Recently there has been an increasing interest in the diet-gut-kidney axis, whereby elements derived from the diet alter the composition of the gut microbiota and production of microbial metabolites which induce effects at extra-intestinal sites, including the kidneys [2, 3]. It has been noted that patients with diabetes [4] and end stage renal disease [5, 6] have a contraction in the bacterial taxa that produce beneficial short chain fatty acids (SCFAs). Furthermore, during end stage renal disease, there is an increase in intestinal permeability and subsequent inflammation [7]. SCFAs act via local metabolite-sensing receptors in order to reduce intestinal permeability and inflammation [8, 9]. The use of dietary therapies that directly target the gut microbiota to increase SCFA production, including probiotics and prebiotics, have been recently considered as potential adjunct interventions to limit injury in diabetic nephropathy [10].

Butyrate acts as a ligand for the G-protein-coupled-receptor (GPR)109a, decreasing intestinal inflammation and promoting gut epithelial barrier integrity, thus GPR109a activation is considered to be protective [11]. A recent study showed that GPR109a modulated the renoprotective effect of butyrate on adriamycin-induced nephropathy [3], however, the role of GPR109a in the development of diabetic nephropathy has not been determined.

This study aimed to assess the effects of insufficient GPR109a signalling via genetic deletion of GPR109a on the development of renal injury in diabetic nephropathy. *Gpr109a*−/− mice or their wildtype littermates (*Gpr109a+/+*) were rendered diabetic with streptozotocin (STZ). Mice received a control diet or an isocaloric, high fiber diet containing 12.5% resistant starch for 24 weeks and gastrointestinal permeability and renal injury were determined.

## Materials and Methods

### Animals

Male mice homozygous for a deletion in the GPR109a receptor (*Gpr109a*−/−), were obtained from Professor Charles Mackay (Monash University, Victoria, Australia) [11], and crossbred with wildtype (WT) C57BL6/J mice purchased from The Jackson Laboratory to produce heterozygous mice, which were then mated to produce *Gpr109a*−/− knockout (KO) mice and littermate WT controls. Mice were housed in a climate-controlled animal facility that had a fixed 12-hour light and 12-hour dark cycle and provided with *ad libitum* access to water and chow. All study protocols were conducted in accordance to the principles and guidelines devised by the Alfred Medical Research & Education Precinct Animal Ethics Committee (AMREP AEC) under the guidelines laid down by the National Health and Medical Research Council (NHMRC) of Australia and had been approved by the AMREP AEC (E1487/2014/B).

### Induction of Diabetes

Diabetes was induced at six weeks of age by five daily intraperitoneal injections of streptozotocin (55 mg/kg Sigma Aldrich) in sodium citrate buffer. Diabetes was confirmed by a glycated haemoglobin (GHb) greater than 8%. Two mice failed to meet this cutoff and were excluded from any further analysis. Of those diabetic mice that were included in the analysis, the mean GHb was 11.7% (median 11.9%).

### Diet Intervention

Since previous studies exploring the role of resistant starch (RS) on the development of renal injury have used supraphysiological doses over a short period of time, we sought to supplement a dose of resistant starch that could be more reasonably expected to be consumed by people (25% HAMS Hi maize 1043, equivalent to 12.5% RS) over a longer time frame (24 weeks). From six weeks of age, mice received either a custom-made control diet (CON) or a high fiber diet supplemented with resistant starch prepared by Speciality Feeds (Perth, Western Australia, Australia). Both of these semi-pure diets were formulated based on a modified AIN93G growth diet for rodents. These diets were isocaloric, had equivalent protein, provided as 20% g/g casein, and fat, provided as 7% g/g canola oil. Each diet contained 5% g/g sucrose, 13.2% g/g dextrinised starch and 7.4% g/g cellulose. A resistant starch supplemented diet (SF15-015) was formulated with 25% g/g Hi-maize 1043, whilst the CON diet (SF15-021) contained an additional 20% g/g regular starch and 5% g/g cellulose in order to maintain caloric equivalency between diets. Hi-maize 1043, an RS2 starch prepared from high amylose maize starch (HAMS) which contains 50% resistant starch [12], was provided as a raw ingredient by Ingredion (Westchester, IL, USA). Mice received these experimental diets *ad libitum* for 24 weeks.

### Tissue collection

At the end of the study period, mice were anaesthetised by an intraperitoneal injection of 100 mg/kg body weight sodium pentobarbitone (Euthatal; Sigma-Aldrich, Castle Hill, Australia) followed by cardiac exsanguination. Following cardiac exsanguination, blood was immediately centrifuged at 6000 rpm for 6 minutes and plasma was snap frozen on dry ice and stored at −80°C. Kidney sections were fixed in neutral buffered formalin (10% v/v). The gastrointestinal tract was dissected and the mesentery removed. Sections of the gastrointestinal tract were weighed and length measured. The ileum was flushed with chilled phosphate buffered saline. Ileum sections were fixed in paraformaldehyde (4% v/v) for 24 hours before being transferred to 4% sucrose solution and embedded in paraffin. Ileum sections were snap frozen in liquid nitrogen and stored at −80°C, for ribonucleic acid (RNA) analysis.

### Glycated haemoglobin

Glycated haemoglobin (GHb) was measured in blood collected at cull using a Cobas b 101 POC system (Roche Diagnostics, Forrenstrasse, Switzerland) according to the manufacturer’s instructions. The Cobas b 101 POC system has a detection range of between 4-14%, with any sample with a GHb less than 4% designated as low and samples with a GHb of greater than 14% designated high.

### *In Vivo* Intestinal Permeability Assay

Intestinal permeability was assessed *in vivo* using the previously described dextran FITC technique [13], during the week prior to cull. In brief, mice were fasted for a minimum of four hours and received an oral gavage of a 125 mg/mL solution of dextran FITC equivalent to 500 mg/kg body weight. After one hour, approximately 120 μL was collected from the tail vein using heparinised capillary tubes. Blood was centrifuged at 6000 rpm for 6 minutes, plasma collected and the fluorescence in plasma samples was determined in relation to a standard dilutions set, using a fluorescence spectrophotometer (BMG Labtech, Ortenberg, Germany) set to excitation 490nm, emission 520nm. The intra- and interassay coefficients of variation were 3.2 and 8.9%, respectively.

### Body Composition

Fat mass and lean body mass were determined using a 4-in-1 EchoMRI body composition analyser (Columbus Instruments, Columbus, OH, USA), which measures fat mass, lean mass and total water content using nuclear magnetic resonance relaxometry [14]. The weight of mice prior to being placed in the body composition analyser was used for calculation of percentage fat and lean mass.

### Metabolic Caging, Urine and Plasma Analyses

After 23 weeks of experimental diet, mice were housed individually in metabolic cages (Iffa Credo, L’Arbresle, France) for 24 hours for urine collection and measurement of urine output and food and water intake. The animals received *ad libitum* access to food and water during this period. Urine was stored at −80°C until required for analyses. Urinary albumin was determined using a mouse specific ELISA (Bethyl Laboratories, Montgomery, TX, USA) according to the kit protocol. The intra- and interassay coefficients of variation were 7.3 and 8.9%, respectively. Urinary monocyte chemoattractant protein-1 (MCP-1) was measured using a commercially available ELISA kit (R&D Systems, Minneapolis MN, USA) as per the kit protocol. The intra− and interassay coefficients of variation were 2.8 and 4.0%, respectively. Blood urea nitrogen was analysed using a commercially available colorimetric urea assay (Arbor Assays, Ann Arbor, MI, USA) as per the kit protocol. The intra− and interassay coefficients of variation were 4.5 and 3.9%, respectively. Plasma cystatin C was determined using a commercially available ELISA from R&D Systems. The intra− and interassay coefficients of variation were 3.8 and 8.3%, respectively

### Kidney and Ileum Histology

Kidneys were fixed in 10% (v/v) neutral buffered formalin prior to embedding in paraffin. Kidney sections (3 μm) were stained with periodic acid-Schiff (PAS) and assessed in a semiquantitative manner, whereby a blinded researcher assessed the level of glomerulosclerosis for each glomerulus and assigned an integer score of between 1 and 4, indicative of the level of severity of glomerulosclerosis. Twenty-five glomeruli were scored per animal, and these scores were averaged to provide a glomerulosclerosis score index (GSI) for each animal, as previously described [15]. Ileal sections were fixed in 4% paraformaldehylde for 24 hours, followed by a transfer to 4% sucrose, and subsequent embedding in paraffin. Ileal sections (5 μm) were stained with haemotoxylin and eosin (H&E) and images were captured using a brightfield microscope (Nikon Eclipse-Ci; Nikon, Tokyo, Japan) coupled with a digital camera (Nikon DS-Fi3; Nikon, Tokyo, Japan). Morphological measurements of villus height and crypt depth were conducted using ImageJ (Version 1.52a). Villus height was measured from the topmost point of the villus to the crypt transition, whilst the crypt depth was measured as the invagination between two villi to the basement membrane.

### Quantitative RT-PCR

RNA was isolated from snap frozen ileum sections using a phenol-chloroform extraction method and used to synthesise cDNA, as previously described [15]. Gene expression of zonulin and occludin was determined using TaqMan (Life Technologies) and SYBR Green reagents (Applied Biosystems), respectively. Gene expression was normalised to 18S mRNA utilising the ΔΔCt method and reported as fold change compared to WT non-diabetic mice receiving the control diet.

### Statistical Analyses

Data were analysed by two-way ANOVA with the Tukey posthoc test for multiple comparisons. Analyses were performed using GraphPad Prism Version 7.01 (GraphPad Software, La Jolle, CA, USA). Data are shown as mean ± SEM. A value of p<0.05 was considered statistically significant.

## Results

### Metabolic and Phenotypic parameters

Consistent with the diabetic phenotype, mice with STZ-induced diabetes had increased glycated haemoglobin and decreased body weight (Table 1). Diabetes was associated with a decrease in relative fat mass and an increased lean mass and increased 24-hour urine output and water intake (Table 1). Neither deletion of the GPR109a receptor nor consumption of the high fiber (resistant starch) diet was associated with changes in glycated haemoglobin, body weight or body composition. Diabetes was associated with an increased liver weight and a decreased spleen weight (Table 1). Mice with diabetes had an increase in small intestine length (p<0.001, Fig 1A) caecum length (p<0.0001, Fig 1B), small intestine weight (p<0.0001, Fig 1D), caecum weight (p<0.0001, Fig 1E) and colon weight (p<0.0001, Fig 1F). Consumption of the fiber supplement led to an increase in caecal weight and length in diabetic mice, but not in non-diabetic mice (Fig 1B, 1E). Resistant starch supplementation was also associated with an increase in colon weight (Fig 1F).

**Figure 1:**
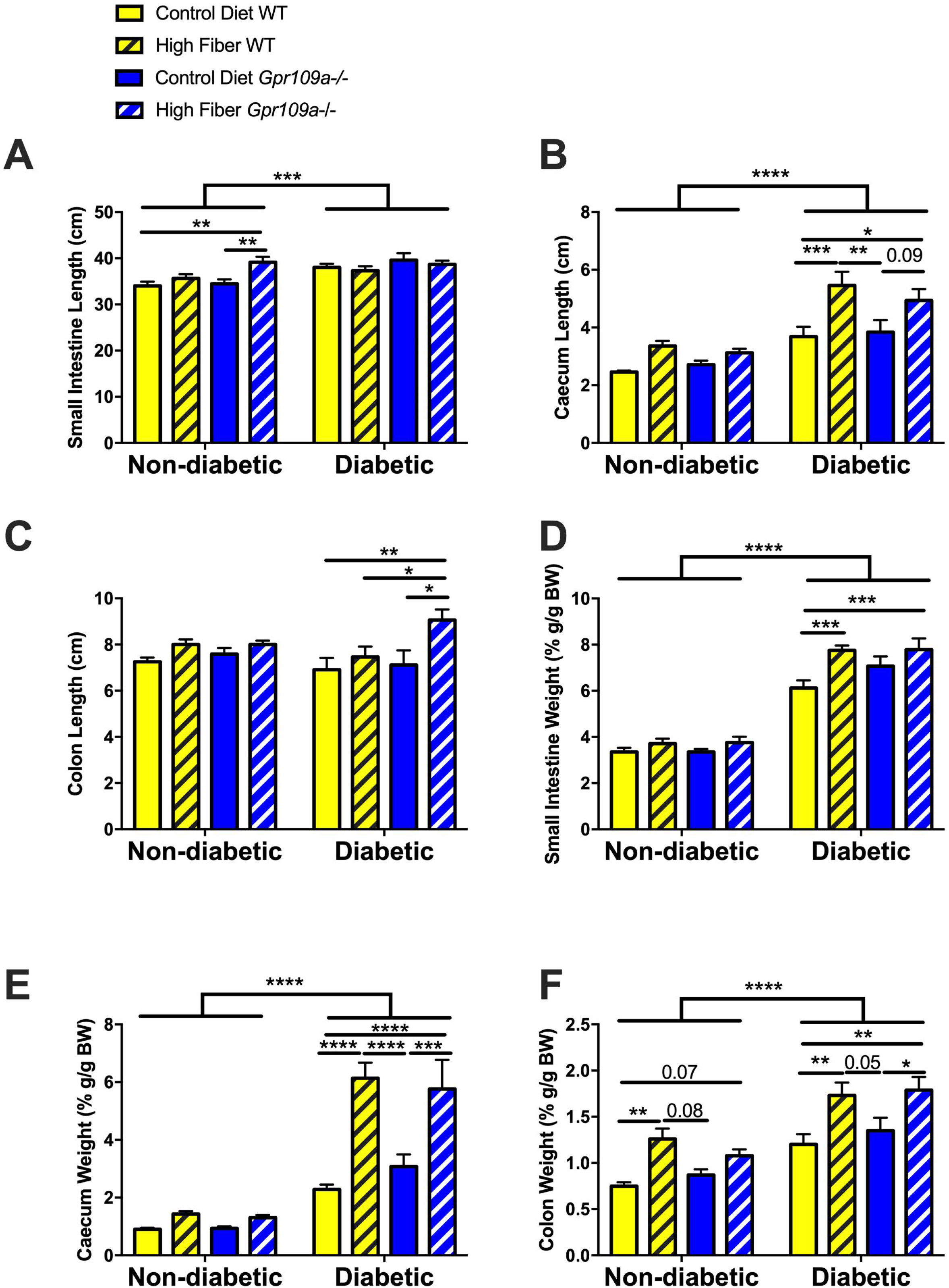
Intestinal Anatomy.

**A** Small intestine length, **B** Caecum length, **C** Colon length, **D** Small intestine weight, **E** Caecum weight, **F** Colon weight. Data are expressed as mean ± S.E.M. * p<0.05, ** p<0.01, *** p<0.001, **** p<0.0001. n = 7-12. WT = wildtype.

**Table 1:**
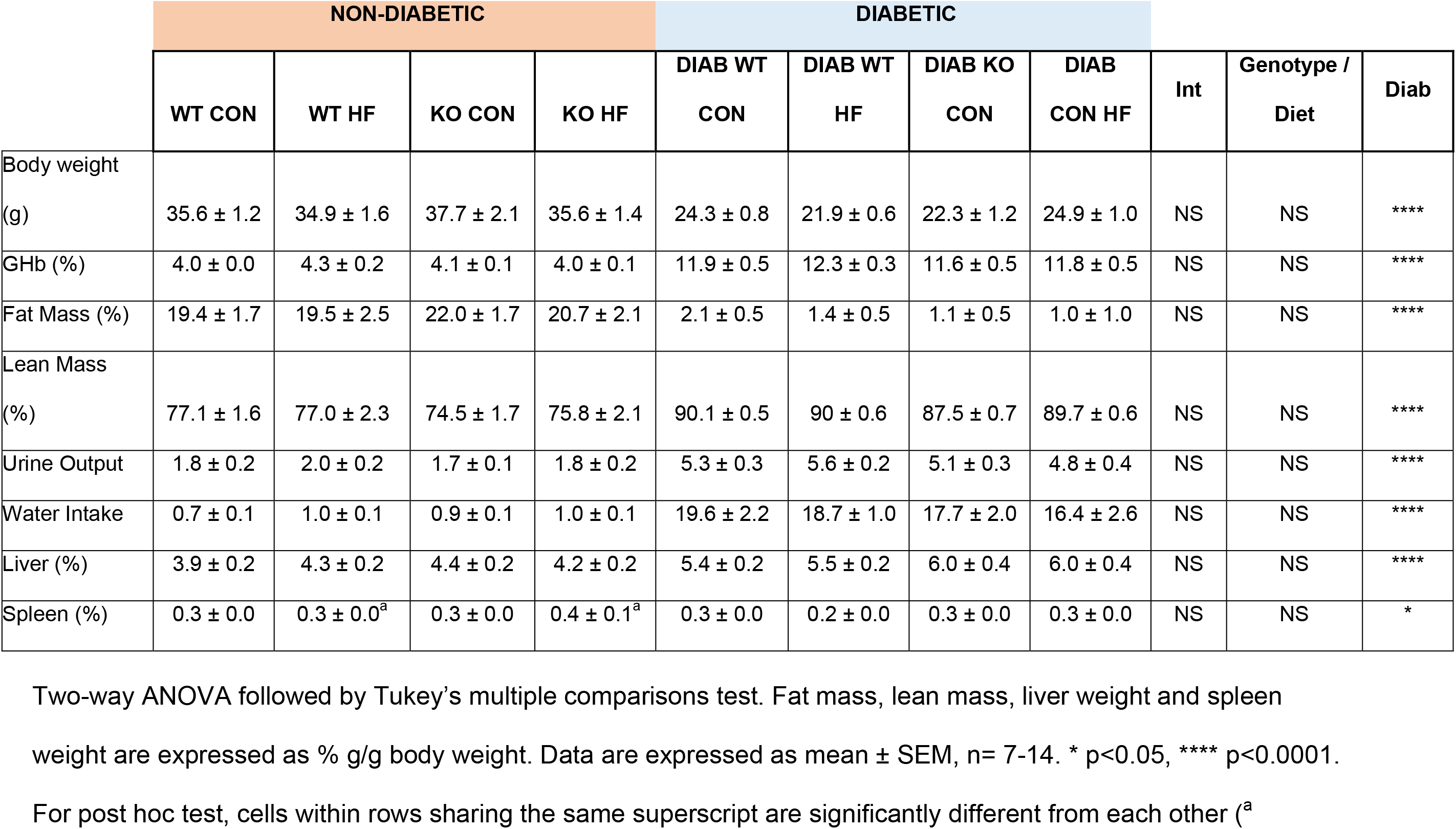
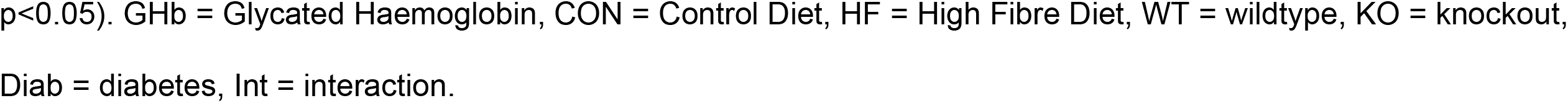
Phenotypic and Biochemical Characteristics of Mice.

|  | NON-DIABETIC |  |  |  | DIABETIC |  |  |  | Int | Genotype /<br>Diet | Diab |
| --- | --- | --- | --- | --- | --- | --- | --- | --- | --- | --- | --- |
|  | WT CON | WT HF | KO CON | KO HF | DIAB WT<br>CON | DIAB WT<br>HF | DIAB KO<br>CON | DIAB<br>CON HF |  |  |  |
| Body weight<br>(g) | 35.6 ± 1.2 | 34.9 ± 1.6 | 37.7 ± 2.1 | 35.6 ± 1.4 | 24.3 ± 0.8 | 21.9 ± 0.6 | 22.3 ± 1.2 | 24.9 ± 1.0 | NS | NS | **** |
| GHb (%) | 4.0 ± 0.0 | 4.3 ± 0.2 | 4.1 ± 0.1 | 4.0 ± 0.1 | 11.9 ± 0.5 | 12.3 ± 0.3 | 11.6 ± 0.5 | 11.8 ± 0.5 | NS | NS | **** |
| Fat Mass (%) | 19.4 ± 1.7 | 19.5 ± 2.5 | 22.0 ± 1.7 | 20.7 ± 2.1 | 2.1 ± 0.5 | 1.4 ± 0.5 | 1.1 ± 0.5 | 1.0 ± 1.0 | NS | NS | **** |
| Lean Mass<br>(%) | 77.1 ± 1.6 | 77.0 ± 2.3 | 74.5 ± 1.7 | 75.8 ± 2.1 | 90.1 ± 0.5 | 90 ± 0.6 | 87.5 ± 0.7 | 89.7 ± 0.6 | NS | NS | **** |
| Urine Output | 1.8 ± 0.2 | 2.0 ± 0.2 | 1.7 ± 0.1 | 1.8 ± 0.2 | 5.3 ± 0.3 | 5.6 ± 0.2 | 5.1 ± 0.3 | 4.8 ± 0.4 | NS | NS | **** |
| Water Intake | 0.7 ± 0.1 | 1.0 ± 0.1 | 0.9 ± 0.1 | 1.0 ± 0.1 | 19.6 ± 2.2 | 18.7 ± 1.0 | 17.7 ± 2.0 | 16.4 ± 2.6 | NS | NS | **** |
| Liver (%) | 3.9 ± 0.2 | 4.3 ± 0.2 | 4.4 ± 0.2 | 4.2 ± 0.2 | 5.4 ± 0.2 | 5.5 ± 0.2 | 6.0 ± 0.4 | 6.0 ± 0.4 | NS | NS | **** |
| Spleen (%) | 0.3 ± 0.0 | 0.3 ± 0.0<sup>a</sup> | 0.3 ± 0.0 | 0.4 ± 0.1<sup>a</sup> | 0.3 ± 0.0 | 0.2 ± 0.0 | 0.3 ± 0.0 | 0.3 ± 0.0 | NS | NS | * |

Two-way ANOVA followed by Tukey's multiple comparisons test. Fat mass, lean mass, liver weight and spleen weight are expressed as % g/g body weight. Data are expressed as mean ± SEM, n= 7-14. \* p<0.05, \*\*\*\* p<0.0001. For post hoc test, cells within rows sharing the same superscript are significantly different from each other (<sup>a</sup>)
p<0.05). GHb = Glycated Haemoglobin, CON = Control Diet, HF = High Fibre Diet, WT = wildtype, KO = knockout, Diab = diabetes, Int = interaction.

### Renal Injury and Inflammation

Diabetic mice exhibited the hallmarks of diabetic nephropathy, including albuminuria (p<0.0001, Fig 2A), increased blood urea nitrogen (p<0.0001, Fig 2B), renal hypertrophy (p<0.0001, Fig 2C) and hyperfiltration (p<0.0001, Fig 2D). Genetic ablation of *Gpr109a* resulted in no change in the renal phenotype in diabetic mice (Fig 2A-D). Likewise, the high fiber diet in the context of diabetes did not alter hallmarks of diabetic nephropathy (Fig 2A-D). Diabetes was associated with an increase in the inflammatory marker MCP-1, whilst there was no effect of either resistant starch supplementation or deletion of *Gpr109a* (p<0.0001, Fig 2E). Assessment of renal histology revealed an overall increase in glomerulosclerosis in the diabetic setting, however, genetic ablation of *GPR109a* or supplementation with the high fiber diet did not impact on renal structural injury (p<0.0001, Figure 3A & B).

**Figure 2:**
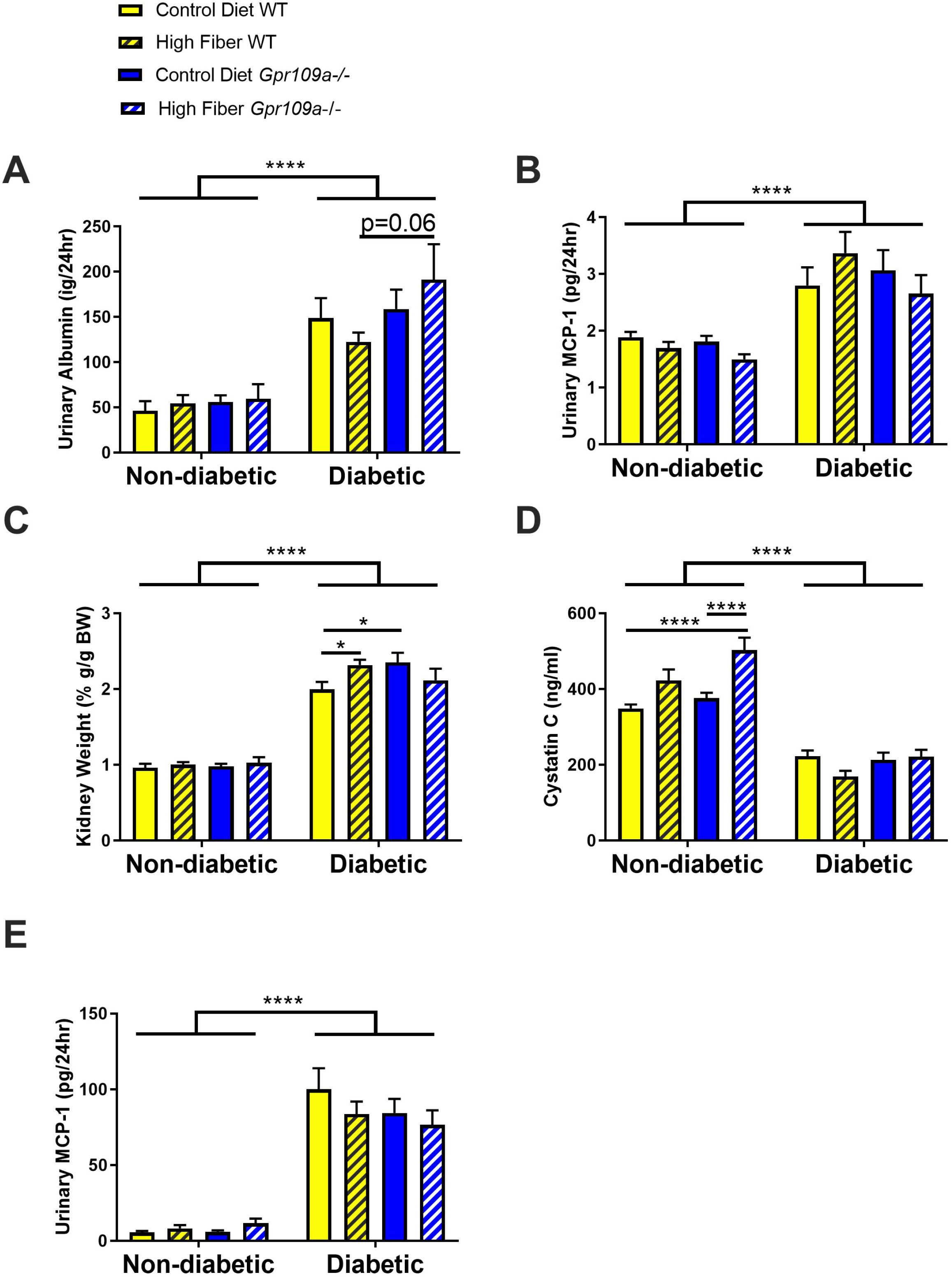
Renal Injury and Inflammation. **A** Urine albumin, **B** Blood urea nitrogen, **C** Relative kidney weight, **D** Plasma Cystatin C, **E** Urinary MCP-1. Data are expressed as mean ± S.E.M. * p<0.05, **** p<0.0001. n = 7-14. MCP-1 = Monocyte Chemoattractant Protein-1.

**Figure 3:**
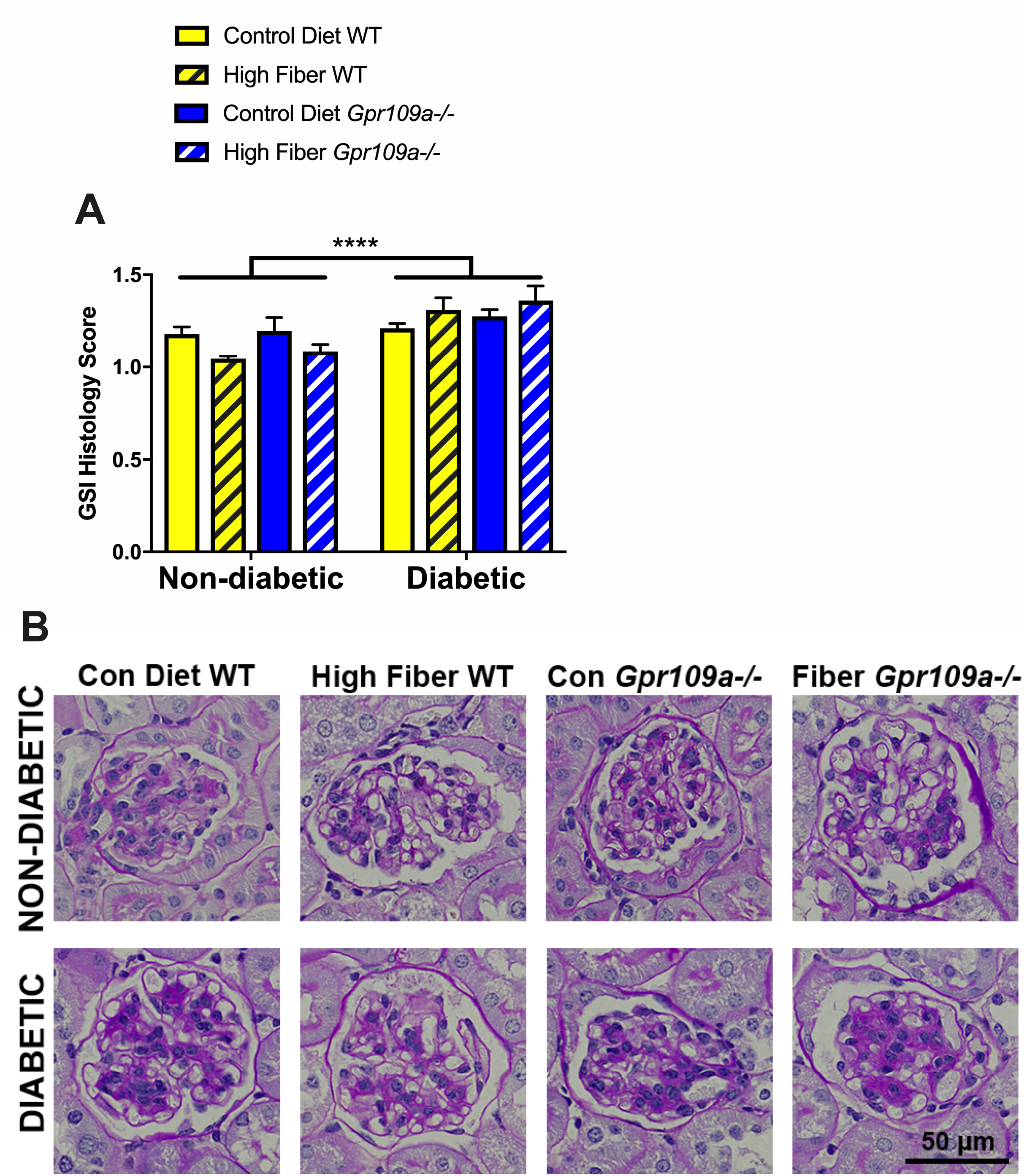
Renal Structural Changes. **A** Glomerulosclerotic Index Score, **B** Representative PAS stained images. Data are expressed as mean ± S.E.M. **** p<0.0001. n = 7-12.

### Intestinal Permeability and Morphology

To determine whether *Gpr109a* deletion led to changes in intestinal morphology, histology of the ileum was undertaken. Diabetes was associated with an overall increase in villi height (p<0.0001, Fig 4A & C) and a reduction in crypt depth (p<0.001, Fig 4B & C) in the ileum. In non-diabetic mice, deletion of *GPR109a* was associated with a trend towards increased villi height (p=0.06, Fig 4A). The high fiber diet led to a trend towards an increase in villi height in non-diabetic wildtype and in *Gpr109a*−/− diabetic mice (Fig 4A). Despite these morphological changes in the ileum with diabetes, diabetes did not induce any alteration in intestinal permeability, as measured by an *in vivo* intestinal permeability procedure (dextran-FITC, Fig 4D) or as determined by gene expression of the tight junction proteins occludin (*Ocln*, Fig 4E) or zonulin (*Tjp-1*) (Fig 4F).

**Figure 4:**
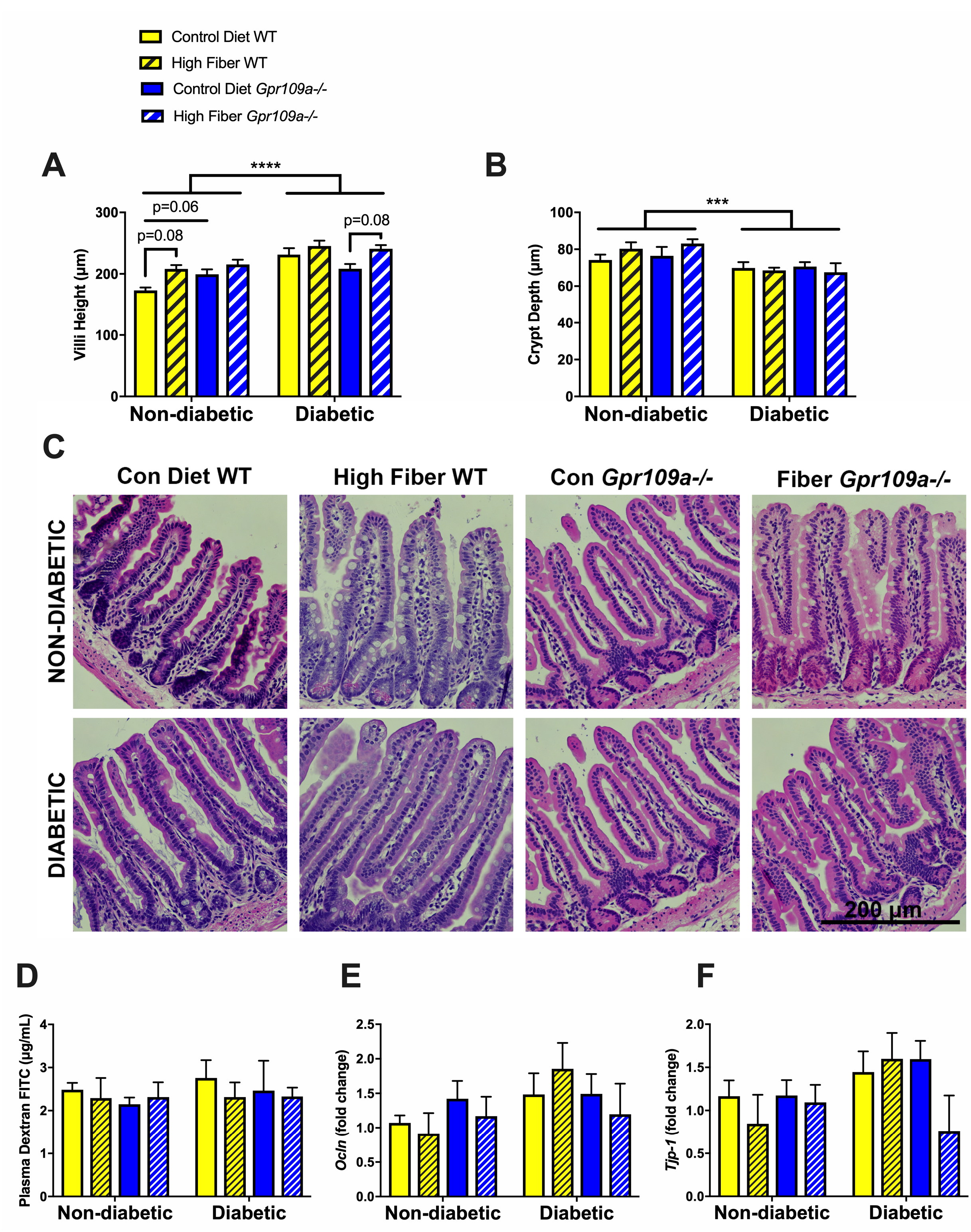
Intestinal Permeability and Morphology. **A** Ileum villi height, **B** Ileum crypt depth, **C** Representative H&E stained images, **D** Plasma dextran-FITC intestinal permeability assay, **E** Ileum expression of Occludin (*Ocln*), **F** Ileum expression of Zonula occludens (*Tjp-1*). Data are expressed as mean ± S.E.M. *** p<0.001, **** p<0.0001. n= 7-12.

## Discussion

The current study explored the effects genetic deletion of the butyrate receptor, GPR109a, had on the development of chronic renal injury in diabetic nephropathy. No effect on the diabetic renal phenotype was observed with deletion of *Gpr109a*. Surprisingly, there was no change in intestinal permeability or morphometry as a result of *Gpr109a* deletion. Furthermore, a high fiber 12.5% resistant starch diet was not effective in reducing renal injury in the setting of diabetes.

This is the first study to assess the effects of deletion of GPR109a on diabetic nephropathy and these findings show that deletion of this receptor was not associated with any change in renal injury in long term studies (24 weeks). Given the hypothesis that renal injury would occur downstream of alterations in intestinal permeability and there was no effect of *Gpr109a* deletion on *in vivo* assessment of intestinal permeability, this should perhaps not come as a surprise finding. *In vitro*, *Gpr109a* knockdown inhibits butyrate-induced increases in the tight junction protein Claudin-3, suggesting that GPR109a may have a role in the integrity of the intestinal epithelial barrier [9]. However *in vivo*, whilst deletion of GPR109a was associated with a trend towards an increase in intestinal permeability in an induced food allergy model [16] there was no effect in otherwise healthy mice [8], indicating that deletion of the GPR109a receptor alone is insufficient to alter intestinal permeability. There is redundancy in the metabolite-sensing GPCR family, with butyrate being recognised by GPR109a, GPR41 and GPR43 receptors. Indeed, it has been suggested that given this redundancy, single knockout models are insufficient to fully elucidate the effects of these receptors, and that double or triple knockout models are required [17].

Resistant starch is a type of dietary fibre that acts as a prebiotic. Numerous studies have illustrated that supplementation with resistant starch is associated with an increase in the microbial production of SCFAs, particularly butyrate [10]. A commonly used source of resistant starch is high amylose maize starch (HAMS) and in an obese model of diabetes, the Zucker diabetic fatty rat, six weeks of supplementation with a diet containing 55% HAMS (Amylogel, equivalent to 20% resistant starch) was associated with a reduction in albuminuria [18]. In addition, in the adenine-induced rat model of chronic kidney disease, supplementation of 59% HAMS diet (Hi-maize 260, equivalent to 27% resistant starch) for three weeks was associated with improvements in creatinine clearance [19]. Conversely however, a study that supplemented a diet containing 55% HAMS (Amylogel, equivalent to 20% resistant starch) for four weeks in male Sprague-Dawley rats with streptozotocin-induced diabetes did not find any renoprotective benefit with resistant starch supplementation [20]. Whilst these results show promise, the concentrations of HAMS used in these studies are likely to be much greater than could be reasonably expected to be consumed by people.

Since previous studies exploring the role of resistant starch on the development of renal injury have used supraphysiological doses over a short period of time, we sought to supplement a dose of resistant starch that could be more reasonably expected to be consumed by people (25% HAMS Hi maize 1043, equivalent to 12.5% resistant starch) over a longer time frame (24 weeks). In the present study, we investigated the effects of resistant starch supplementation, at a dose of ~12.5%, on the development of diabetic nephropathy in the STZ-induced diabetic mouse. Supplementation of a high fiber resistant starch diet in such chronic studies was not renoprotective.

A recent study showed that a renoprotective effect of butyrate on adriamycin-induced nephropathy in short term studies [3]. In that study, intraperitoneal injection of butyrate for 7 or 14 days in a model of adriamycin-induced nephropathy; and dietary supplementation of a high fiber butyrylated resistant starch diet for four weeks prior to adriamycin-induced nephropathy (preventative approach), showed that butyrate led to a decreased urinary protein to creatinine ratio. Although no measures of intestinal permeability were determined, that study shows promise for butyrate as a renoprotective agent, at least in the acute setting, most likely attributable to acute inflammation.

In conclusion, this study shows that long term resistant starch supplementation at a dosage that would reasonably be expected to be consumed by humans, did not alleviate albuminuria in the STZ-induced diabetes model. Whilst diabetes was associated with alterations in intestinal morphology, there was no change in intestinal permeability. It would be pertinent to consider resistant starch supplementation in other rodent models of diabetes that may be more representative of the intestinal changes that occur with diabetes in humans. Finally, this study indicates that GPR109a deletion does not play a critical role in the development of diabetic nephropathy nor gastrointestinal homeostasis.

## Acknowledgements

The authors would like to thank the following people: Maryann Arnstein for technical assistance, Professor Charles R. Mackay (Monash University) for the *GPR109a*−/− mice, Warren Potts from Specialty Feeds, Australia for the design and generation of the diets. MS and RSJL were supported by scholarships from the Australian government Research Training Program. GCH was supported by a postdoctoral fellowship from JDRF. SMT is supported by a JDRF Advanced Postdoctoral Fellowship and MTC is supported by a Career Development Award from the JDRF Type 1 Diabetes Clinical Research Network, a special research initiative of the Australian Research Council.

